# Isolation and comparative analysis of antibodies that broadly neutralize sarbecoviruses

**DOI:** 10.1101/2021.12.11.472236

**Authors:** Lihong Liu, Sho Iketani, Yicheng Guo, Ryan G. Casner, Eswar R. Reddem, Manoj S. Nair, Jian Yu, Jasper F-W. Chan, Maple Wang, Gabriele Cerutti, Zhiteng Li, Candace D. Castagna, Laura Corredor, Hin Chu, Shuofeng Yuan, Vincent Kwok-Man Poon, Chris Chun-Sing Chan, Zhiwei Chen, Yang Luo, Marcus Cunningham, Alejandro Chavez, Michael T. Yin, David S. Perlin, Moriya Tsuji, Kwok-Yung Yuen, Peter D. Kwong, Zizhang Sheng, Yaoxing Huang, Lawrence Shapiro, David D. Ho

**Affiliations:** Aaron Diamond AIDS Research Center, Columbia University Vagelos College of Physicians and Surgeons, New York, NY 10032, USA; Department of Microbiology and Immunology, Columbia University Vagelos College of Physicians and Surgeons, New York, NY 10032, USA; Zuckerman Mind Brain Behavior Institute, Columbia University, New York, NY 10027, USA; Department of Biochemistry and Molecular Biophysics, Columbia University Vagelos College of Physicians and Surgeons, New York, NY 10032, USA; State Key Laboratory of Emerging Infectious Diseases, Carol Yu Centre for Infection, Department of Microbiology, Li Ka Shing Faculty of Medicine, The University of Hong Kong, Hong Kong Special Administrative Region, China; Centre for Virology, Vaccinology and Therapeutics, Health@InnoHK, Hong Kong Special Administrative Region, China; Institute of Comparative Medicine, Columbia University Irving Medical Center, New York, NY 10032, USA; AIDS Institute, Li Ka Shing Faculty of Medicine, The University of Hong Kong, Hong Kong Special Administrative Region, China; Hackensack Meridian Health Center for Discovery and Innovation, Nutley, NJ 07110, USA; Hackensack Meridian School of Medicine, Nutley, NJ 07110, USA; Department of Pathology and Cell Biology, Columbia University Vagelos College of Physicians and Surgeons, New York, NY 10032, USA; Division of Infectious Diseases, Department of Medicine, Columbia University Vagelos College of Physicians and Surgeons, New York, NY 10032, USA; Vaccine Research Center, National Institutes of Health, Bethesda, MD 20892, USA

## Abstract

The devastation caused by SARS-CoV-2 has made clear the importance of pandemic preparedness. To address future zoonotic outbreaks due to related viruses in the sarbecovirus subgenus, we identified a human monoclonal antibody, 10-40, that neutralized or bound all sarbecoviruses tested *in vitro* and protected against SARS-CoV-2 and SARS-CoV *in vivo*. Comparative studies with other receptor-binding domain (RBD)-directed antibodies showed 10-40 to have the greatest breadth against sarbecoviruses and thus its promise as an agent for pandemic preparedness. Moreover, structural analyses on 10-40 and similar antibodies not only defined an epitope cluster in the inner face of the RBD that is well conserved among sarbecoviruses, but also uncovered a new antibody class with a common CDRH3 motif. Our analyses also suggested that elicitation of this class of antibodies may not be overly difficult, an observation that bodes well for the development of a pan-sarbecovirus vaccine.

**One sentence summary:** A monoclonal antibody that neutralizes or binds all sarbecoviruses tested and represents a reproducible antibody class.

## Main Text

The COVID-19 pandemic is caused by the severe acute respiratory syndrome coronavirus 2 (SARS-CoV-2) that has infected >245 million people and resulted in >5 million deaths (*1*). Multiple variants of this virus have emerged, including some capable of increased transmission or antibody evasion (*2, 3*). Furthermore, the threat of continued zoonotic spillovers warrants the development of interventions that could broadly combat animal coronaviruses with pandemic potential (*4*).

Numerous anti-SARS-CoV-2 monoclonal antibodies (mAbs) have been isolated and characterized, with several demonstrating clinical utility (*5, 6*). Some have been reported to possess broadly neutralizing activity against not only SARS-CoV-2 but also other sarbecoviruses (*7–15*), a viral subgenus containing both SARS-CoV-2 and SARS-CoV (*16*). Such mAbs could serve as a therapeutic adjunct for the current pandemic as well as a useful agent in addressing future zoonoses due to a sarbecovirus. We now report the isolation of three mAbs that broadly neutralize sarbecoviruses. In addition, we describe virological and structural findings from a comprehensive comparative analysis of these mAbs together with those previously reported to have broad activity. The information provided herein could aid the development of pan-sarbecovirus antibodies and vaccines.

To isolate mAbs with the desired neutralization breadth, we screened sera from convalescent COVID-19 patients for neutralizing activity against a panel of variant viruses. Serum from Patient 10 and Patient 11 potently neutralized all SARS-CoV-2 variants tested as well as SARS-CoV, albeit weakly (**Fig. S1**). We then sorted for B.1.351 spike trimer-specific memory B cells from the blood of both patients, followed by single-cell RNA-sequencing to determine the paired heavy and light chain sequences of each mAb (**Fig. S2**) (*17*). A total of 58 mAbs were isolated and characterized.

Three mAbs, 10-40, 10-28, and 11-11, were found to bind to SARS-CoV-2 spikes of variants D614G and B.1.351 as well as the SARS-CoV spike (**Fig. 1A**). All three antibodies recognized epitopes within the receptor binding domain (RBD) (**Fig. 1A**) and inhibited the binding of soluble human ACE2 receptor to the spike (**Fig. 1B**). Epitope mapping by competition ELISA was carried out on these 3 antibodies along with a panel of 9 RBD-specific mAbs reported to have breadth against sarbecoviruses, including DH1047 (*7*), S2X259 (*8*), REGN10985 (*9*), ADG-2 (*10*), 2-36 (*11*), COVA1-16 (*12*), CR3022 (*13*), S2H97 (*14*), and S309, also known as sotrovimab (*15*). 10-40, 10-28, and 11-11 fell into one competition group with 7 other mAbs (**Fig. 1C and S3**) that are known to recognize an inner face of RBD when it is in the “up” position (*7–12*). The epitope of S2H97 was partially overlapping whereas that of S309 was discrete, not surprisingly since the latter is directed to an epitope on the outer face on RBD (*14, 15*). The binding affinities of this panel of mAbs to SARS-CoV-2 and SARS-CoV spikes were measured by surface plasmon resonance and summarized in **Fig. S4**.

**Fig. 1.**
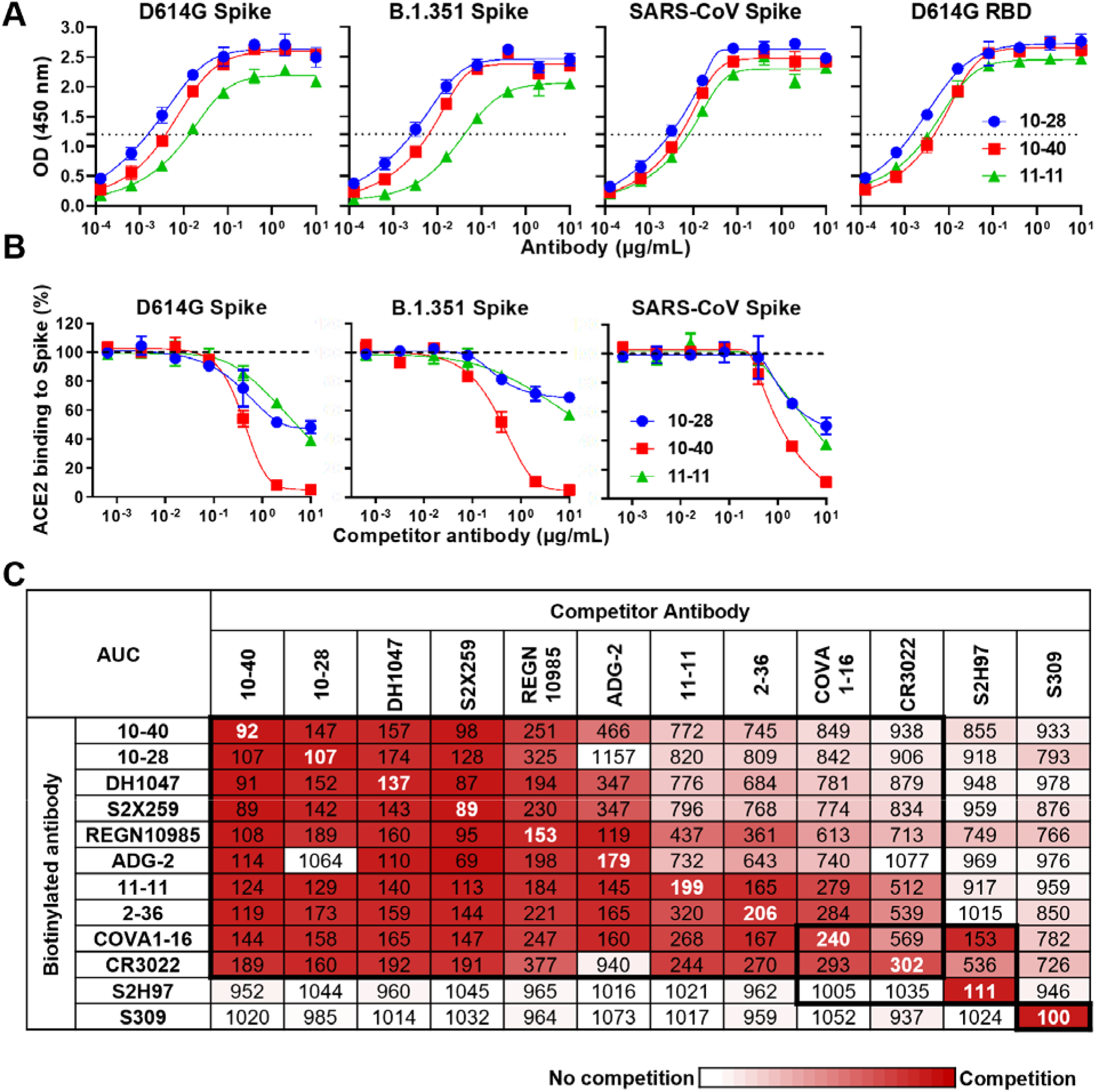
Identification of three mAbs that bind to the spikes of SARS-CoV-2 D614G, B.1.351, SARS-CoV, as well as the RBD of D614G. **(A) and inhibit ACE2 binding to SARS-CoV-2 D614G, B.1.351, and SARS-CoV spikes (B)**. Data are shown as mean ± SD of two technical replicates. **(C)** Epitope mapping by competition ELISA of newly identified mAbs together with other RBD-directed broadly neutralizing mAbs. A representative result of three experimental replicates is shown.

Genetically, 10-40, 10-28, and 11-11 utilized IGHV4-39*01, IGHV3-30*18, and IGHV4-31*03 heavy-chain V genes with CDRH3 lengths of 22, 13, and 21 amino acids, respectively. The light chains of 10-40, 10-28, and 11-11 were derived from IGLV6-57*01, IGKV1-39*01, and IGLV1-40*01, respectively (**Fig. S5A**). All three antibodies had low levels of somatic hypermutation (**Fig. S5, A and B**).

We then comprehensively compared the virus-neutralizing potency and breadth of 10-40, 10-28, and 11-11 to other RBD-directed mAbs with known activity against other sarbecoviruses. First, each antibody was evaluated against SARS-CoV-2 variants in neutralization assays using both VSVΔG-pseudotyped viruses and authentic viruses. All mAbs, except CR3022, showed breadth by neutralizing all SARS-CoV-2 strains tested (**Figs. 2A, 2B, S6, and S7A**). ADG-2 was the most potent, followed by a group comprised of 10-40, DH1047, S2X259, REGN10985, and S309. In general, 10-28, 11-11, 2-36, and COVA1-16 exhibited lower potencies. Against authentic SARS-CoV, ADG-2, 10-40, S2X259, and DH1047 showed decent neutralizing activity with IC50 values well below 1 µg/mL (**Figs. 2B and S7B**).

**Fig. 2.**
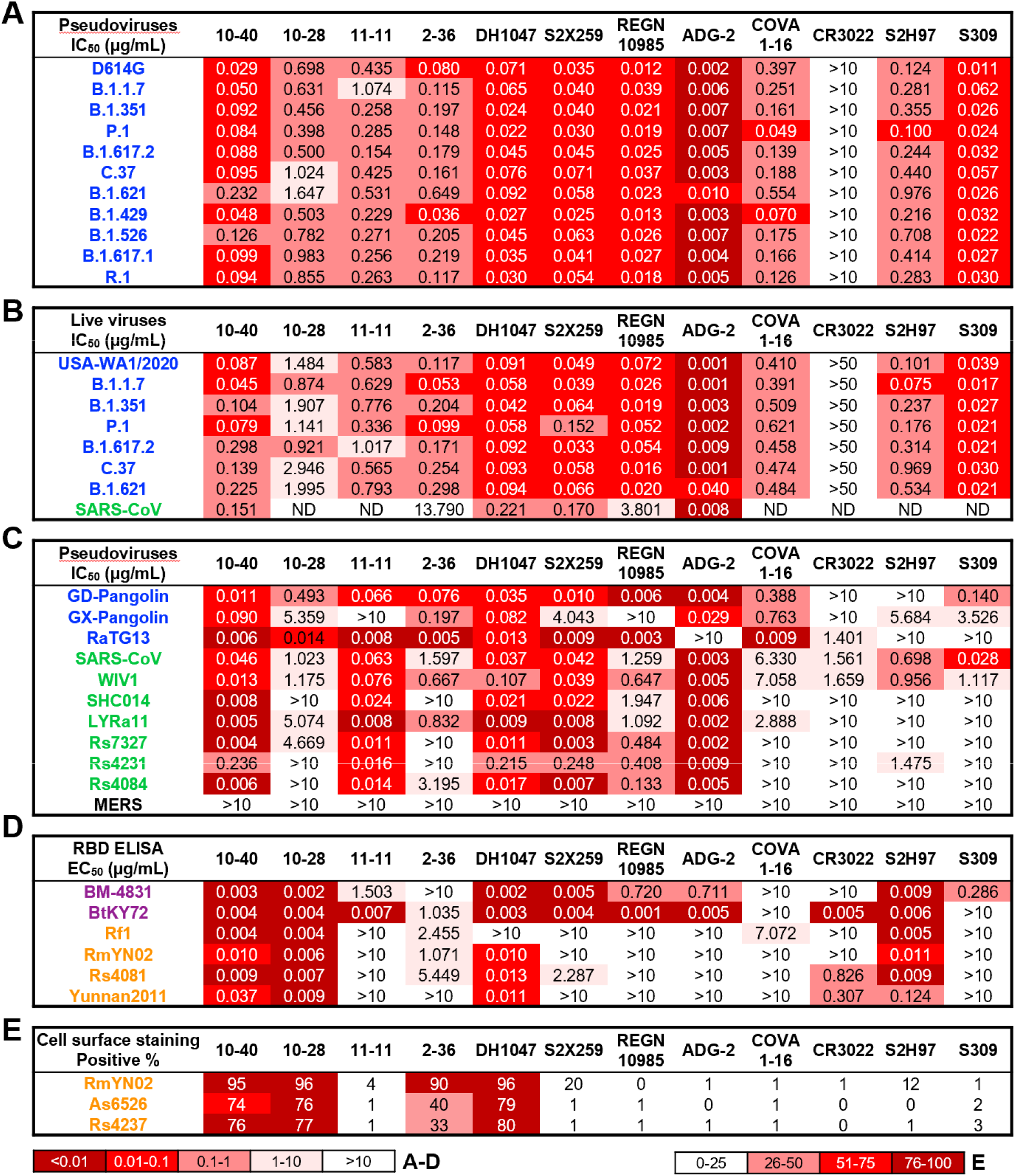
Breadth and potency of 10-40, 10-28, and 11-11 versus other reported antibodies with broad reactivity. Neutralization titers (IC50) against pseudoviruses of SARS-CoV-2 variants **(A)**, authentic SARS-CoV-2 variants and SARS-CoV GZ50 strain **(B)**, and pseudoviruses of other animal sarbecoviruses in the SARS-CoV-2 (blue) and SARS-CoV (green) sublineages **(C)**. Binding of mAbs to purified RBD proteins from African/European (purple) or Asian (orange) bat sarbecoviruses as measured by ELISA (EC50) **(D)**, or to spike proteins expressed on the surface of transfected cells by flowcytometry **(E)**. A representative result of three experimental replicates is shown. ND = not determined.

We next evaluated each mAb for neutralization against ten non-SARS-CoV-2 sarbecoviruses capable of using human ACE2 as receptor in a VSVΔG-pseudotyped virus assay (**Figs. 2C and S8**). Only 10-40 and DH1047 neutralized all sarbecoviruses tested, whereas 11-11, S2X259, and ADG-2 were deficient or weak in neutralizing at least one of the viruses in the panel. The other mAbs had many more “holes” in their repertoire. As expected, none of these antibodies neutralized the Middle East respiratory syndrome virus (MERS), a member of the merbecovirus subgenus.

There are more sarbecoviruses found in bats in Africa/Europe or Asia (**Fig. S9A**) that do not use human ACE2 as receptor (*16, 18*). Since their target cells are unknown, performing virus-neutralization assay is not readily feasible. We therefore examined the binding profiles of this panel of mAbs to RBD proteins derived from six sarbecoviruses outside of SARS-CoV-2 and SARS-CoV sublineages (**Figs. 2D and S9B**). 10-40, 10-28, and S2H97 bound all RBDs tested, whereas DH1047 did not recognize the RBD of Rf1.

The remaining mAbs bound only a subset of the RBDs. In particular, S2X259, REGN10985, ADG-2, and S309 did not recognize, largely, the RBD of Asian bat sarbecoviruses. Similarly, we assessed the binding of this panel of mAbs to three spikes of Asian bat sarbecoviruses (**Fig. S9A**) as expressed on the surface of transfected cells. 10-40, 10-28, 2-36, and DH1047 exhibited breadth, but other mAbs were largely non-reactive (**Figs. 2E and S10**).

In summary, the totality of findings in **Fig. 2** shows that 10-40 can neutralize or bind every sarbecovirus we have studied, and it appears to be an RBD-directed mAb with the greatest breadth against sarbecoviruses known to date.

To investigate the nature of antibody-spike interactions for 10-40, 10-28, and 11-11, we determined the cryo-EM structures for Fabs of these mAbs in complex with S2P-prefusion-stabilized spike proteins from SARS-CoV-2 WA-1 and B.1.351 strains. Interestingly, a greater degree of spike disassembly was observed for all WA-1 complexes, and higher quality cryo-EM maps were obtained for all three B.1.351 complexes. A single predominant population was observed where three Fabs were bound per spike in a 3-RBD-up conformation (**Fig. 3A, S11, S12, S13, and Table S1**). For 11-11, an additional class of two Fabs bound with 2-RBD up was also observed. The 10-40 complex reconstruction reached 3.5 Å global resolution, but local refinement of the RBD + Fab maps did not surpass 4 Å resolution. To resolve the interfaces, we determined crystal structures of the 10-40 and 10-28 Fab:RBD complexes, as elaborated in paragraphs below. Both the crystal structures of 10-40 and 10-28 fitted nearly perfectly in the cryoEM reconstruction density (**Fig. S11 and S12**). For 11-11, a homology model was built and docked into the map (**Fig. S13**), which showed this antibody recognizes RBD in the same way as S2X259 (*8*), with both sharing a similar light chain and CDRH3 motif.

**Fig. 3.**
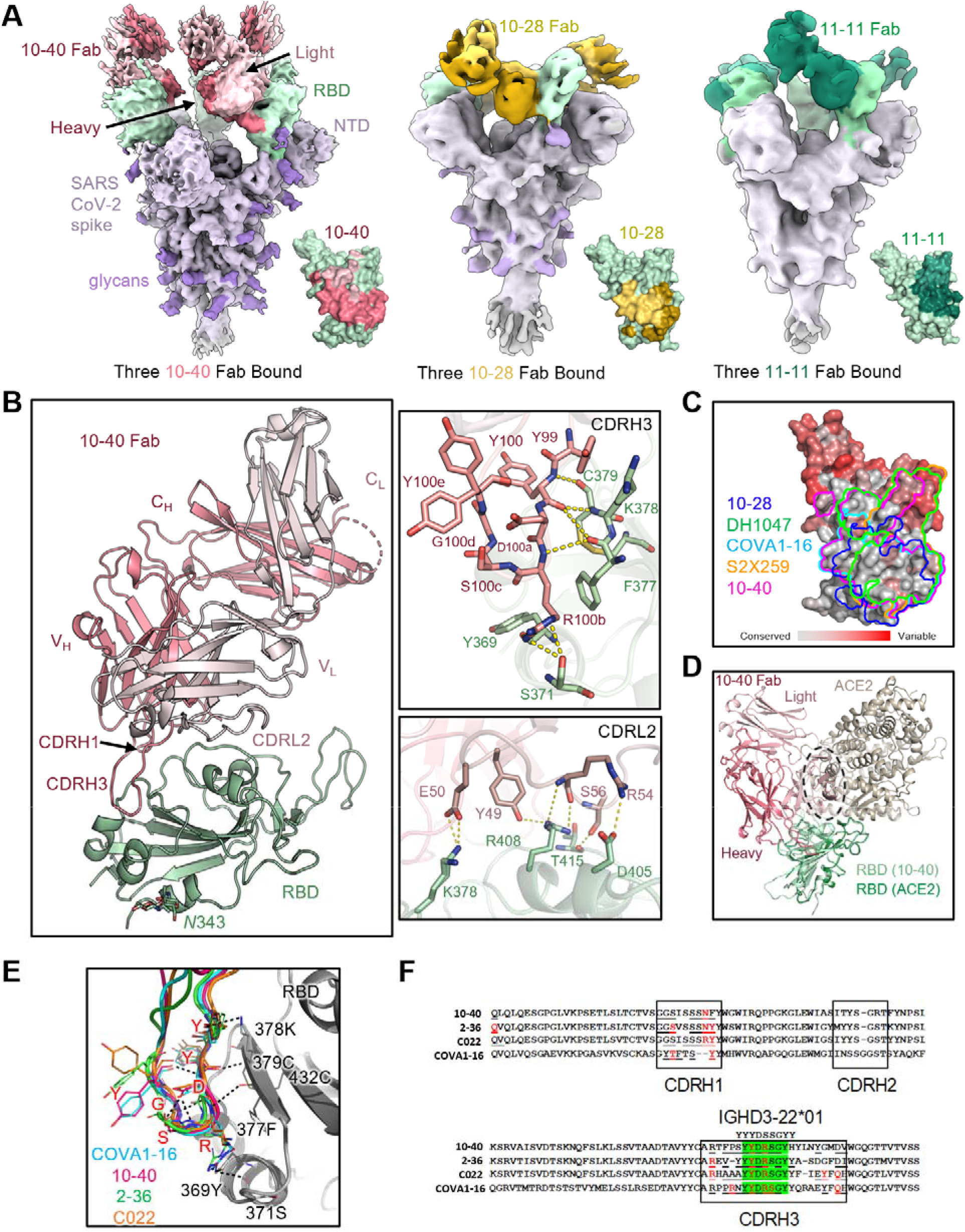
Structural and bioinformatic analyses of isolated antibodies. **(A)**Cryo-EM reconstructions and recognition footprints for 10-40, 10-28, and 11-11 Fabs bound to B.1.351 spike trimer. The spike is colored in light gray, with the RBDs in green and the glycans in purple, oriented with the membrane towards the bottom. The 10-40 Fabs are colored in red, the 10-28 Fabs in yellow, and the 11-11 Fabs in dark green. The Fab heavy chains are shaded darker than the light chains. The footprint of each respective antibody on the inner face of RBD is displayed next to each spike. **(B)** Crystal structure of 10-40 Fab bound to WA1 SARS-CoV-2 RBD. **(C)** Comparison of 10-28, DH1047, COVA1-16, S2X259, and 10-40 epitope footprints on SARS-CoV-2 RBD. The RBD was colored according to the sequence conservation of each residue across 52 sarbecoviruses. **(D)** Overlay of 10-40 and ACE2 binding to RBD, showing a clash between the 10-40 light chain and ACE2. **(E)** Comparison of 10-40, 2-36, C022, and COVA1-16 CDRH3s produced by superimposing RBDs from each complex, revealing a similar binding mode. **(F)** Heavy chain sequence alignment of 10-40, 2-36, C022, and COVA1-16, with the CDRH3 aligned to the IGHD3-22*01 gene. The paratope residues are underlined and residues that form hydrogen bonds with RBD are colored in red.

To visualize the epitopes of 10-40, 10-28, and 11-11 on the SARS-CoV-2 WA1 RBD at higher resolution, we determined the structures of Fab in complex with RBD using X-ray crystallography, with suitable crystals obtained for 10-40 and 10-28 (**Table S2**). The 10-40 crystal structure at 1.5 Å resolution revealed recognition of an epitope that is highly conserved among sarbecoviruses on the inner face of RBD (**Figs. 3B and 3C**). 10-40 used three of its six CDR loops: H1, H3, and L2, to interact with an epitope consisting of a loop on RBD (residues 377-385) and extended toward the RBD ridge near the ACE2 binding site. 10-40 also established extensive polar contacts and hydrophobic interactions with RBD residues (**Fig. 3B**).

For the 10-28 crystal structure (**Fig. S14A**), antibody-RBD side-chain interactions were well defined at 3.2 Å resolution. Interactions were mediated by 10-28 CDR loops H1, H3, L1, and L3, which predominantly contacted α3, but also α2 and the β2-α3, α4-α5 and α5-β4 loops of the RBD. The heavy chain formed three hydrogen bonds and a single salt bridge, between D95 (CDRH3) and K386 (RBD), while the light-chain residues formed a total of five hydrogen bonds (**Fig. S14B**).

All three antibodies recognized a region on the inner side of RBD that is hidden in the RBD-down conformation of spike. Thus, they can only recognize RBD in the up conformation. The 10-40 epitope is similar to the ‘class 4’ antibody epitope previously defined for COVA1-16, C022 and 2-36 (**Fig. S15A**) (*19*). Superposition of the 10-40 Fab with the ACE2-RBD complex (6M0J) showed that antibody binding places the VL domain of the antibody in a position that would clash with ACE2 (**Fig. 3D**), consistent with the experimental data showing inhibition of ACE2 binding (**Fig. 1B**). In addition, these mAbs all targeted the same exposed β-sheet on RBD from a similar angle. S2X259 and DH1047 also recognized the same β-sheet, but from a different angle, similar to 11-11 (**Fig. S15B**). The epitope of 10-28 also included a portion of this β-sheet although the focus was lower on the RBD, similar to CR3022 (**Fig. S15C**).

Given the broad recognition of 10-40 for sarbecoviruses, we carefully analyzed the amino acids that form its epitope and noted their remarkable conservation among sarbecoviruses (**Figs. 3C, S16A, and S16B**), suggesting that this RBD region must have been subjected to strong functional constraints during evolution of this subgenus. Residues 377-379 in a β-sheet was specifically targeted by the CDRH3 of 10-40 via multiple hydrogen bonds (**Fig. 3E**). Interestingly, COVA1-16, 2-36, and C022 interacted quite similarly with the same residues. We also noticed that these three mAbs, together with 10-40, contact this particular β-sheet through a ‘YYDRSGY’ motif originating from IGHD3-22 (**Fig. 3F**), a D gene that is frequently used by antibodies in the human repertoire (**Fig. S17**). This motif contains a Ser-to-Arg substitution, which formed hydrogen bonds with 369Y and 371S on RBD (**Fig. 3E**). Importantly, we believe these structural similarities define 10-40, 2-36, C022, and COVA1-16 as members of a new antibody class, each using a shared mode of heavy-chain binding to RBDs of sarbecoviruses (**Fig. 3F**). As these four mAbs use diverse heavy-chain V genes and light-chain recombinations (**Fig. S18**) and show low-level somatic hypermutation (**Fig. S5**), the elicitation of this class of antibodies may not be overly difficult. This observation bodes well for the development of a pan-sarbecovirus vaccine.

Finally, we evaluated the *in vivo* protective efficacy of 10-40 by challenging wild-type mice with a mouse-adapted SARS-CoV-2 strain, MA10 (*20*), or hACE2-transgenic mice with SARS-CoV (*21*). The RBD mutations in MA10 (Q493K, Q498Y, and P499T) mapped outside of the 10-40 epitope (**Fig. S19A**) and did not strongly affect the neutralizing activity of 10-40 *in vitro* (**Fig. S19B**). We then performed a prevention experiment (**Fig. 4A**), administering 10-40 or an anti-HIV-1 control mAb 24 hours before the mice were challenged intranasally with MA10. Compared to the control group, significant weight loss was prevented (10 mg/kg) or mitigated (2 mg/kg) by 10-40 administration (**Fig. 4B**). In mice given the control antibody, high levels of infectious virus were observed in the lungs (>10^5^ TCID50/g lung), while little (≤10^4^ TCID50/g lung) or no infectious virus was found in mice given 10-40 at 2 mg/kg or 10 mg/kg, respectively (**Fig. 4C**). An analogous prevention experiment was conducted against SARS-CoV in hACE2-transgenic mice (**Fig. 4D**). Weight loss was again prevented by 10-40 administration (**Fig. 4E**), and levels of SARS-CoV were markedly reduced in the lungs of mice pre-treated with 10-40 (**Fig. 4F**). 10-40 appeared to be active *in vivo* against both SARS-CoV-2 and SARS-CoV.

**Fig. 4.**
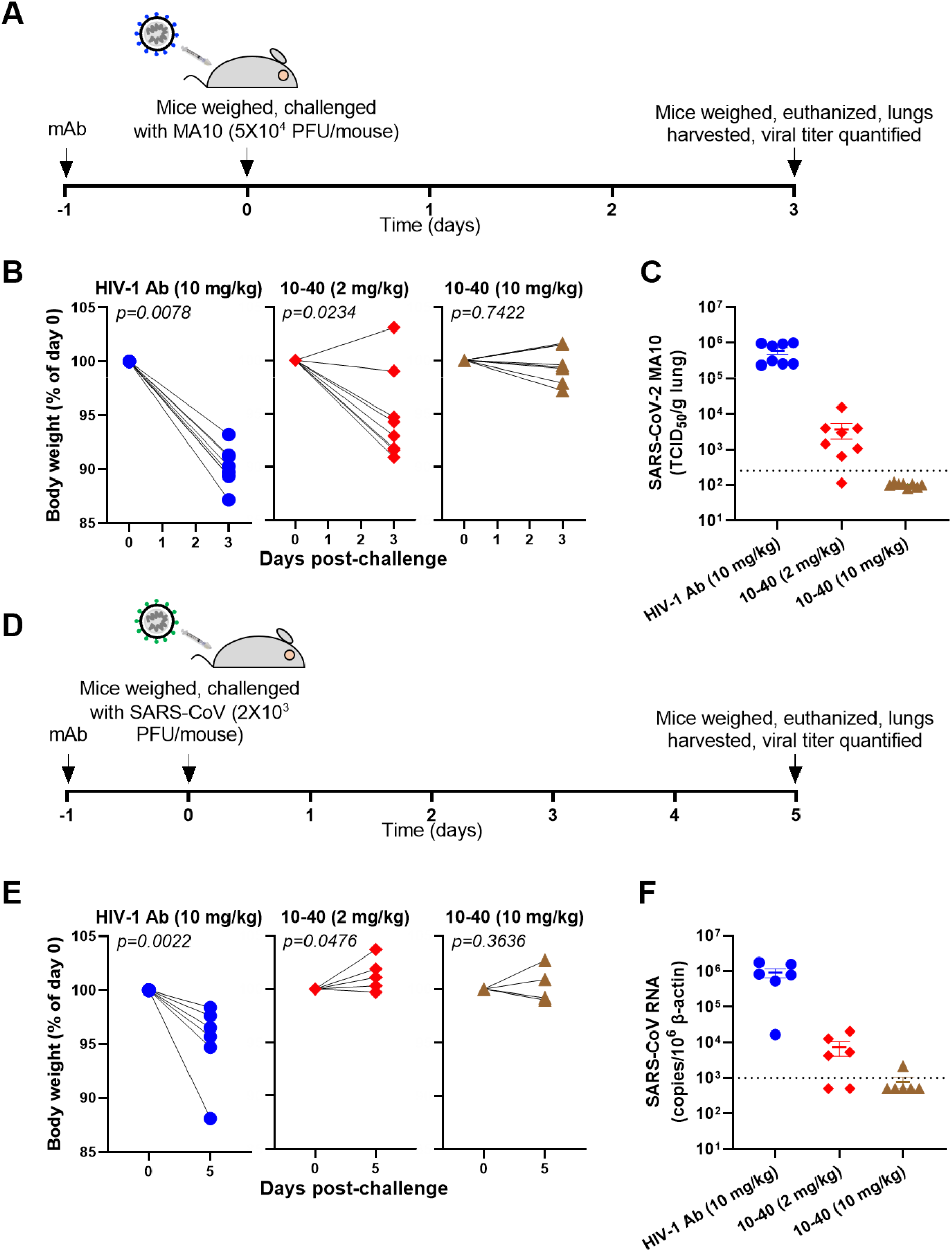
Prophylactic protection against a mouse-adapted strain of SARS-CoV-2 (MA10) and SARS-CoV by 10-40. **(A)** Experimental timeline of the protection study in MA10-challenged mice. **(B)** Body weight change of individual mice in each treatment group (n=8 mice/group). P values were determined by two-tailed t-test with Wilcoxon matched-pairing. **(C)** TCID50/g of lung from individual mice in each treatment group. **(D)** Experimental timeline of the protection study in SARS-CoV-challenged mice. **(E)** Body weight change of individual mice in each treatment group (n=6 mice/group). P values were determined by two-tailed t-test with Wilcoxon matched-pairing. **(F)** SARS-CoV RNA (normalized to β-actin) within lung from individual mice.

Sarbecoviruses have caused two major outbreaks in humans in the past two decades: SARS-CoV in 2002-2003 and SARS-CoV-2 now. Surely, the world must prepare for the possibility of a future epidemic/pandemic due to another member of this subgenus that is presently harbored by bats and other animals (*4, 16, 22*). A pan-sarbecovirus neutralizing mAb and/or vaccine could be useful interventions to contain another outbreak. We have identified a human mAb, 10-40, that suits this need. While RBD-directed mAbs that neutralize sarbecoviruses have been reported (*7–15*), 10-40 shows the greatest breadth in our comprehensive comparative analysis, followed closely by DH1047 (*7*) (**Fig. 2**). 10-40 neutralizes or binds to every sarbecovirus we have tested, regardless of their usage of ACE2 as receptor. It has the requisite potency against SARS-CoV-2 *in vitro* and *in vivo* (**Fig. 4**); interestingly, its potency against other sarbecoviruses is even better (**Fig. 2**), despite being isolated from a COVID-19 patient. Very recently, three sarbecoviruses closest genetically to SARS-CoV-2 have been identified in bats in Laos (*22*), along with two more in a different sublineage (**Fig. S16**). While not empirically tested, we note that they are likely to be susceptible to 10-40 because the key amino acids that would form this RBD epitope are identical to those in either GD-Pangolin or RmYN02 (**Fig. S16**), both of which are neutralized or bound by 10-40 (**Figs. 2C-E**). We believe 10-40 is a promising candidate for pandemic preparedness.

The pursuit of a pan-sarbecovirus vaccine is already underway, including strategies that specifically target the stem helix in the S2 region of spike (*23*) or conserved elements on RBD. Efforts directed to the latter have already shown promise (*24–26*). The structural differences in the epitopes recognized by 10-40 and DH1047 (**Figs. 3C and S15**) versus epitopes recognized by mAbs (such as ADG-2 or S2X259) with lower breadth against sarbecoviruses (**Fig. S15**) could be informative in focusing the antibody response to certain conserved residues on the inner face of RBD. The epitopes of 10-40 and DH1047 are substantially overlapping, but their angles of approach are different (**Fig. S15**), suggesting that this site could be attacked in more than one way. Moreover, by genetically comparing 10-40 with other antibodies, we have uncovered a unique class of mAbs that target a particular β-sheet in RBD through a common CDRH3 motif (**Fig. 3E-F**). That this class of mAbs could use multiple heavy and light chain V genes (**Fig. S18**) and without extensive somatic hypermutation (**Fig. S5**) is certainly welcome news for the development of a pan-sarbecovirus vaccine targeting the RBD.

## Acknowledgments

We are grateful to A. Cohen and P. Bjorkman (Caltech) for RBD constructs, B. Zhang (NIAID) for DH1047, C. Kyratsous (Regeneron Pharmaceuticals) for REGN10985, and L.Y. Liu and H. Wang (Columbia University) for the B.1.621 spike construct. We thank the Columbia University Single Cell Analysis Core for conducting 10X Genomics sequencing. We thank K. Perry and J. Schuermann for help with synchrotron data collection conducted at the APS NE-CAT 24-ID-E beamline, which is supported by National Institutes of Health (NIH) P41 GM103403; use of NE-CAT at the Advanced Photon Source was supported by the US Department of Energy, Basic Energy Sciences, Office of Science, under contract number W-31-109-Eng-38. We thank B. Grassucci, C. Wang, and J. Wang for help with cryo-EM data collection at the Columbia University Cryo-EM Facility at the Zuckerman Institute. This study was supported by funding from the Gates Foundation, JPB Foundation, Andrew and Peggy Cherng, Samuel Yin, Carol Ludwig, David and Roger Wu, and Health@InnoHK.

## Author contributions

D.D.H. conceived this project. L.L. and S.I. sorted antigen-specific memory B cells. Y.G. and Z.S. analyzed the 10X Genomics sequencing data and antibody repertoire. J.Y. and M.W. cloned and expressed antibodies. L.L. and Z.L. expressed and purified sarbecovirus proteins. L.L. conducted ELISA binding, surface plasmon resonance, cell surface staining, and competition experiments. L.L. and S.I. performed pseudovirus neutralization experiments. S.I. and A.C. constructed sarbecovirus spike plasmids.

M.S.N. and Y.H. performed authentic SARS-CoV-2 neutralization experiments. J.F-W.C., H.C., Z.C., and K-Y.Y. performed authentic SARS-CoV neutralization experiments. R.G.C. solved the cryo-EM structures. E.R.R. solved the X-ray crystallography structures. Y.G., R.G.C., E.R.R., G.C., Z.S., and L.S. conducted the structural analyses. M.S.N., C.D.C., L.C., M.T., and Y.H. conducted the SARS-CoV-2 in vivo experiments. J.F-W.C., S.Y., V.K-M.P., C.C-S.C., and K-Y.Y. conducted the SARS-CoV in vivo experiments. Y.L. helped with project management. M.T.Y. and D.S.P. provided clinical samples. P.D.K. contributed to discussions. L.L., S.I., Y.G., R.G.C., E.R.R., M.S.N., J.Y., J.F-W.C., M.T., Z.S., Y.H., L.S., and D.D.H. analyzed the results and wrote the manuscript.

## References

1. E. Dong, H. Du, L. Gardner, An interactive web-based dashboard to track COVID-19 in real time. Lancet Infect Dis 20, 533–534 (2020).

2. P. Wang et al., Antibody resistance of SARS-CoV-2 variants B.1.351 and B.1.1.7. Nature 593, 130–135 (2021).

3. N. G. Davies et al., Estimated transmissibility and impact of SARS-CoV-2 lineage B.1.1.7 in England. Science 372, (2021).

4. S. Simpson, M. C. Kaufmann, V. Glozman, A. Chakrabarti, Disease X: accelerating the development of medical countermeasures for the next pandemic. Lancet Infect Dis 20, e108–e115 (2020).

5. M. Dougan et al., Bamlanivimab plus Etesevimab in Mild or Moderate Covid-19. N Engl J Med 385, 1382–1392 (2021).

6. D. M. Weinreich et al., REGN-COV2, a Neutralizing Antibody Cocktail, in Outpatients with Covid-19. N Engl J Med 384, 238–251 (2021).

7. D. Li et al., In vitro and in vivo functions of SARS-CoV-2 infection-enhancing and neutralizing antibodies. Cell 184, 4203–4219 e4232 (2021).

8. M. A. Tortorici et al., Broad sarbecovirus neutralization by a human monoclonal antibody. Nature 597, 103–108 (2021).

9. R. Copin et al., The monoclonal antibody combination REGEN-COV protects against SARS-CoV-2 mutational escape in preclinical and human studies. Cell 184, 3949–3961 e3911 (2021).

10. C. G. Rappazzo et al., Broad and potent activity against SARS-like viruses by an engineered human monoclonal antibody. Science 371, 823–829 (2021).

11. P. Wang et al., A monoclonal antibody that neutralizes SARS-CoV-2 variants, SARS-CoV, and other sarbecoviruses. bioRxiv, (2021).

12. H. Liu et al., Cross-Neutralization of a SARS-CoV-2 Antibody to a Functionally Conserved Site Is Mediated by Avidity. Immunity 53, 1272–1280 e1275 (2020).

13. M. Yuan et al., A highly conserved cryptic epitope in the receptor binding domains of SARS-CoV-2 and SARS-CoV. Science 368, 630–633 (2020).

14. T. N. Starr et al., SARS-CoV-2 RBD antibodies that maximize breadth and resistance to escape. Nature 597, 97–102 (2021).

15. D. Pinto et al., Cross-neutralization of SARS-CoV-2 by a human monoclonal SARS-CoV antibody. Nature 583, 290–295 (2020).

16. M. F. Boni et al., Evolutionary origins of the SARS-CoV-2 sarbecovirus lineage responsible for the COVID-19 pandemic. Nat Microbiol 5, 1408–1417 (2020).

17. L. Liu et al., Potent neutralizing antibodies against multiple epitopes on SARS-CoV-2 spike. Nature 584, 450–456 (2020).

18. H. L. Wells et al., The evolutionary history of ACE2 usage within the coronavirus subgenus Sarbecovirus. Virus Evol 7, veab007 (2021).

19. C. O. Barnes et al., SARS-CoV-2 neutralizing antibody structures inform therapeutic strategies. Nature 588, 682–687 (2020).

20. S. R. Leist et al., A Mouse-Adapted SARS-CoV-2 Induces Acute Lung Injury and Mortality in Standard Laboratory Mice. Cell 183, 1070–1085 e1012 (2020).

21. P. B. McCray, Jr. et al., Lethal infection of K18-hACE2 mice infected with severe acute respiratory syndrome coronavirus. J Virol 81, 813–821 (2007).

22. S. Temmam et al., Coronaviruses with a SARS-CoV-2-like receptor-binding domain allowing ACE2-mediated entry into human cells isolated from bats of Indochinese peninsula. Res Sq, (2021).

23. J. T. Ladner et al., Epitope-resolved profiling of the SARS-CoV-2 antibody response identifies cross-reactivity with endemic human coronaviruses. Cell Rep Med 2, 100189 (2021).

24. A. A. Cohen et al., Mosaic nanoparticles elicit cross-reactive immune responses to zoonotic coronaviruses in mice. Science 371, 735–741 (2021).

25. D. R. Martinez et al., Chimeric spike mRNA vaccines protect against Sarbecovirus challenge in mice. Science 373, 991–998 (2021).

26. A. C. Walls et al., Elicitation of broadly protective sarbecovirus immunity by receptor-binding domain nanoparticle vaccines. Cell 184, 5432–5447 e5416 (2021).

